# Transcriptome analysis of differential gene expression in *Dichomitus squalens* during interspecific mycelial interactions and the potential link with laccase induction

**DOI:** 10.1101/359646

**Authors:** Zixuan Zhong, Nannan Li, Binghui He, Yasuo Igarashi, Feng Luo

## Abstract

Interspecific mycelial interactions between white rot fungi are always accompanied by increased production of laccase. In this study, the potential of white rot fungi *Dichomitus squalens* for enhancing laccase production during interaction with two other white rot fungi *Trametes versicolor* or *Pleurotus ostreatus* was identified. To probe the mechanism of laccase induction and the role of laccase played during the combative interaction, we analyzed the laccase induction response to stressful conditions during fungal interaction related to the differential gene expression profile. We further confirmed the expression patterns of 16 selected genes by qRT-PCR analysis. We noted that many differential expression genes (DEGs) encoding proteins were involved in xenobiotics detoxification and ROS generation or reduction, including aldo/keto reductase, glutathione S-transferases, cytochrome P450 enzymes, alcohol oxidases and dehydrogenase, manganese peroxidase and laccase. Furthermore, many DEG-encoding proteins were involved in antagonistic mechanisms of nutrient acquisition and antifungal properties, including glycoside hydrolase, glucanase, chitinase and terpenoid synthases. DEGs analysis effectively revealed that laccase induction was likely caused by protective responses to oxidative stress and nutrient competition during fungal interspecific interaction.

---

Interspecific interactions between wood-decaying fungi are always associated with competition for territory and resources to support fungal community development (Boddy, 2000; Heilmann-Clausen and Boddy, 2005; Wells and Boddy, 2002). When niches of different species overlap, a broad array of antagonistic responses can be triggered in the interaction zone where mycelia contact occurs. These responses, including changes in mycelial morphology, production of extracellular enzymes and secretion of secondary metabolites (Boddy, 2000; Eyre et al., 2010), play an important role in intensifying the attack or defense mechanism and nutrient uptake to gain advantage over the competitor (Arfi et al., 2013; Eyre et al., 2010).

It has been reported that a range of chemicals, such as alcohols, aldehydes, ketones, terpenes, aromatic compounds and reactive oxygen species (ROS) are produced under antagonism within interaction mycelia (Arfi et al., 2013; Evans et al., 2008), leading to the up-regulation of many oxidative enzymes (Chi et al., 2007; Ferreira Gregorio et al., 2006; Hiscox et al., 2010). Laccase, an enzyme for oxidizing a variety of phenolic and aromatic compounds, plays a defensive role in reducing the oxidative stress caused by oxygen radicals originating from the reaction of toxic molecules (Li et al., 2011; Yang et al., 2012). Furthermore, laccase also synthesized melanin, which is involved in the absorption of toxic compounds and oxygen radicals to protect hyphae against interspecific oxidative stress (Eisenman and Casadevall, 2012; Eisenman et al., 2007; Nosanchuk JD, 2003). Many studies have shown that laccase plays a defensive role against stressful conditions (Cho et al., 2009; Mayer and Staples, 2002; Piscitelli et al., 2011; Zhu and Williamson, 2004), and laccase activity increased during interactions among many white-rot fungi (Baldrian, 2004; Chi et al., 2007; Ferreira Gregorio et al., 2006; Flores et al., 2009; Hiscox et al., 2010; Kuhar et al., 2015). Thus, laccase might be generally involved in the combative response of different white-rot fungi to interspecific interactions. Moreover, increased laccase activity during mycelial interaction also implied intensive nutrient competition (Hiscox et al., 2010). Based on direct combative interactions between mycelia to defend or occupy resources for their own growth (Boddy, 2000), a series of antagonistic metabolites were up-regulated, especially toxic or antifungal compounds, which were likely induced by oxidative stress (El Ariebi et al., 2016; Peiris et al., 2007).

Our preliminary study showed that the laccase activity could be highly increased during the co-culture of *Dichomitus squalens* with two other white rot fungi *Trametes versicolor* and *Pleurotus ostreatus.* Considering that laccase is the most important oxidative enzyme secreted from white rot fungi, many studies focused on the laccase induction particularly in the mycelial interaction of different fungi (Baldrian, 2004; Flores et al., 2009; Kuhar et al., 2015; Wei et al., 2010), but lacking further exploration on the molecular mechanism of interspecific interaction and laccase induction among different fungi. As oxidative stress can occur in the interaction region of different fungi (Silar, 2005), and fungal laccase can be involved in the defense response to oxidative stress, which is also commonly accompanied by the overexpression of a large amount of other antioxidative enzymes (Jaszek et al., 2006; Yang et al., 2012). Thus, in this study, we performed a transcriptional analysis of gene expression changes in *D. squalens* under the interaction with two wood-decaying fungi *T. versicolor* and *P. ostreatus* to investigate the mechanism of fungal competitive interaction, and focus on the potential role including resistance against oxidative stress that laccase played during the interaction of *D. squalens* with other two competitors based on analysis of transcriptomic level.

## MATERIALS AND METHODS

### Strains and culture

Strains of *T. versicolor* and *P. ostreatus* were from the Biological Resource Center, NITE (NBRC), the strain number is NBRC30388 for *T. versicolor* and NBTC30776 for *P. ostreatus. D. squalens* was from the Deutsche Sammlung von Microorganismen and Zellkulturen (DSMZ), and the strain number is DSMZ9615. Strains were maintained on potato dextrose agar (PDA) slants and stored at 4°C. Before use, the stored fungi were inoculated onto the newly prepared PDA plates at 28°C for pre-culturing.

### Co-culture growth conditions

A paired culture of *D. squalens* with *T. versicolor* or *P. ostreatus* was grown on Sc medium (Flundas and Hibbett, 2012), consisting of 10g/liter glucose,1.5g/liter **L-**asparagine, 0.12mg/liter thiamine dichloride, 0.46g /L KH_2_PO_4_, 1g/L K_2_HPO_4_, 0.5g/L MgSO_4_·7 H_2_O, 5 mg/L FeCl_3_·6 H_2_O, 0.06 mg/L HBO3, 0.04 mg/L (NH4)6Mo7O_24_·4H_2_O, 0.2 mg/L CuSO_4_·5H_2_O, 2 mg/L ZnSO_4_·7H_2_O, 0.1 mg/L MnSO_4_·4H_2_O, 0.4 mg/L CoCl_2_·6H_2_O, 1.2 mg/L Ca(NO_3_)_2_·4H_2_O, 20g/L Agar. Co-culture experiments were inoculated with two plugs (Φ7 mm) of the appropriate fungal strains from pre-culture on opposite sides of a 90 mm petri dish containing Sc agar, and the two plugs were kept at a distance of 5 cm. The pure culture of Ds was set as control. All treatments were made with three replicates. After incubation for 9 days at 28°C, three plugs (3 × Φ5 mm) taken from the plate at isolates and interaction zones were immersed in 0.1 M acetic acid buffer (pH 4.8) for 6 hours at 4°C. The crude enzyme samples were centrifuged at a speed of 12000 rpm/min for 2 min at 4°C for laccase activity determination.

### NBT staining assay

To determine ROS in the interaction zone and single species zone of each plate, the nitroblue tetrazolium (NBT) reduction assay was performed using a 0.3 mM NBT aqueous solution with 0.3 mM NADPH (Lara et al., 2003). Mycelia were flooded with stain solution and incubated at room temperature with gentle rotation. Photographs were taken with a Nikon Coolpix camera to show purple coloration resulting from the reduction of NBT by ROS after 30 min staining.

### RNA extraction, library construction and sequencing

The mycelium from single culture (*D. squalens* growing alone only) and two co-cultures DsTv (interaction zone of *D. squalens* and *T. versicolor*) and DsPo (interaction zone of *D. squalens* and *P. ostreatus*) were used as samples for the RNA extractions, each sample set three biological replicates for RNA-seq analysis. All samples were ground into a powder in liquid nitrogen using a mortar and pestle and extracted using the UNIIQ-10 Trizol Total RNA Kit according to the manufacturer’s instructions. The concentrations and quality of purified nucleic acids were checked using the ND-2000 (NanoDrop Technologies) and **Agilent 2100 Bioanalyzer**, and multiplexed libraries were constructed and sequenced on an Illumina HiSeq4000. All samples displayed a 260/280 ratio greater than 2.0 and RNA integrity numbers (RIN) ≥8.0.

To obtain an overview of the gene expression profiles of *D. squalens* during interaction with *T. versicolor* or *P. ostreatus*, a RNA-seq transcriptome library was prepared following the instructions for the TruSeq^™^ RNA sample preparation Kit from Illumina (San Diego, CA) using 5 μg of total RNA. The mRNA was isolated according to the polyA selection method using oligo(dT) beads and fragmented in fragmentation buffer. The double-stranded cDNA is then synthesized using a SuperScript double-stranded cDNA synthesis kit (Invitrogen, CA) with random hexamer primers (Illumina). The synthesized cDNA is subjected to end-repair, phosphorylation and ‘A’ base addition, according to Illumina’s library construction protocol. Libraries were size-selected for cDNA target fragments of 200-300 bp on 2% Low Range Ultra Agarose and PCR amplified using Phusion DNA polymerase (NEB) for 15 PCR cycles. After quantification by TBS380, a paired-end RNA-seq sequencing library was sequenced with Illumina HiSeq4000 (1 × 51 bp read length). The reference genome of *D. squalens* was download in NCBI (https://www.ncbi.nlm.nih.gov/genome) (Floudas et al., 2012).

### Differential expression analysis and functional enrichment

To identify differential expression genes (DEGs) between co-culture and pure culture samples, the expression level of each gene was calculated according to the fragments per kilobase of exon per million mapped reads (FRKM) method. RSEM (http://deweylab.biostat.wisc.edu/rsem/) (Li and Dewey, 2011) was used to quantify gene and isoform abundances. The R statistical package software EdgeR (Empirical analysis of Digital Gene Expression in R, http://www.bioconductor.org/packages/2.12/bioc/html/edgeR.html) (Robinson et al., 2010) was utilized for differential expression analysis. In addition, functional-enrichment analysis including GO and KEGG were performed to identify which DEGs were significantly enriched in GO terms and metabolic pathways at the Bonferroni-corrected p-value < 0.05 compared to the whole-transcriptome background. GO functional enrichment and KEGG pathway analysis were performed with Goatools (https://github.com/tanghaibao/Goatools) and KOBAS (http://kobas.cbi.pku.edu.cn/home.do) (Xie et al., 2011).

### Gene expression analysis by qRT-PCR

Quantitative real-time PCR was conducted in 96-well plates with a 7500 Real Time PCR System (Applied Biosystems) using a SYBR® Premix Ex TaqTM II Kit (TaKaRa, Japan). Each reaction mixture contained 10-μL volumes: 5 μl SYBR® Premix Ex TaqTM II, 0.2 μl ROX reference dye, 0.4 μl forward primer, 0.4 μl reverse primer, 1 μl cDNA template and 3 μl sterile distilled water. The recommended protocol for PCR was used according to the manual: 95 °C for 30 s, 40 cycles of 95 °C for 5 s and 60 °C for 34 s. Three biological replicates per sample and three technical replicates were conducted for each cDNA template. To normalize the qRT-PCR data, 40S ribosomal protein S24 (DICSQDRAFT_46225), chitinase (DICSQDRAFT_98815) and α-tubulin (DICSQDRAFT_135086) with stable expression in all samples were used as the endogenous reference genes, and data were normalized using the average expression level of the three reference genes. The specific primers of target genes used in this work were listed in Table 1. Quantitative Relative quantification was established using the 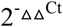 method (Livak and Schmittgen, 2001).

**Table 1.**
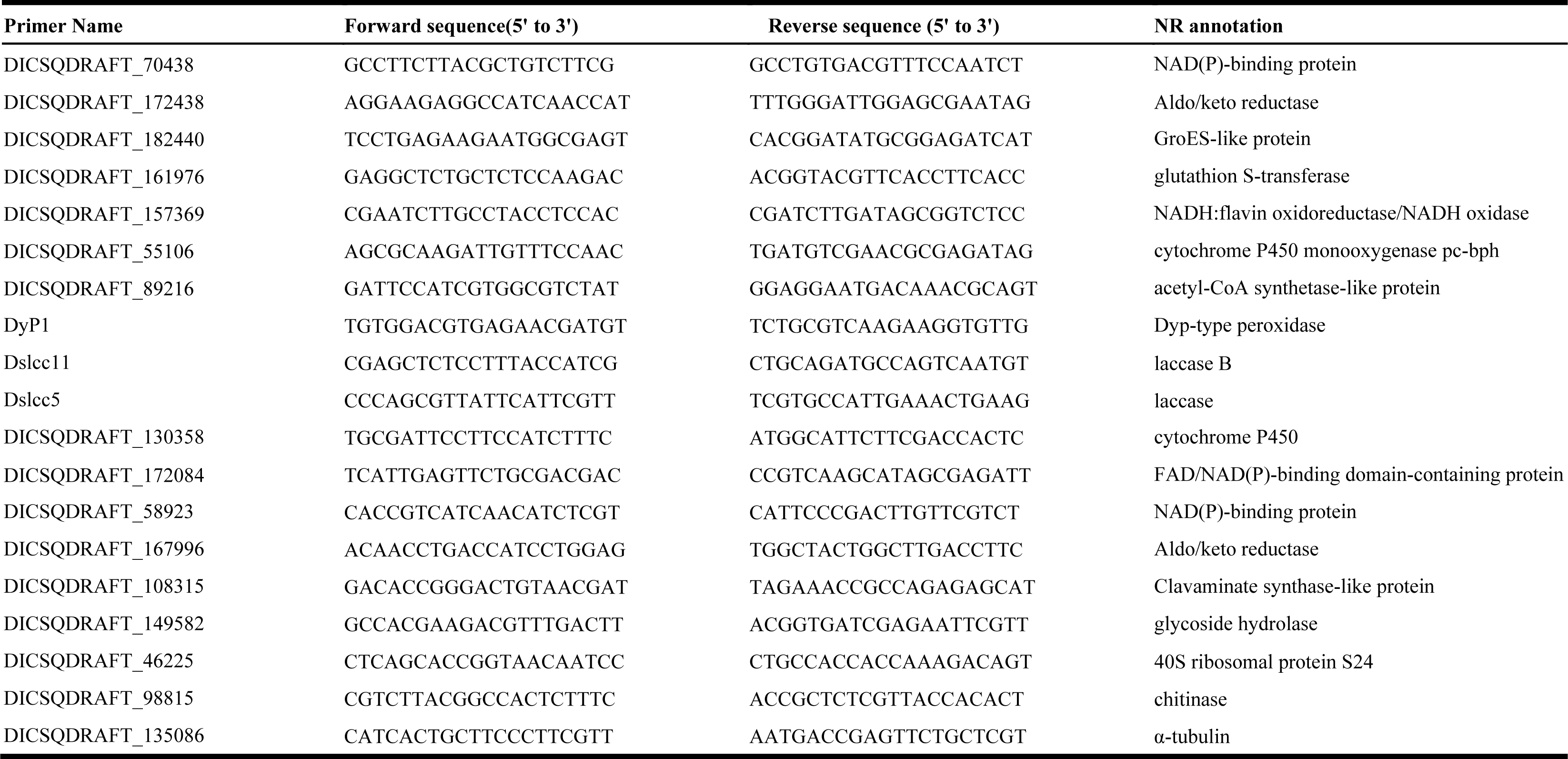
PCR primers used for the q-RT-PCR experiments

### Assays of Laccase

Enzyme assays of crude enzyme samples were conducted in a 3 ml reaction mixtures according to the method described in Eggert et al., 1996 with slight modification (Eggert et al., 1996). Laccase activity was measured spectrophotometrically by monitoring the oxidation of 500 mM ABTS [2,20-azinobis-(3-ethylbenzthiazoline-6-sulphonate)] (molar extinction coefficient 36,000 M^−1^cm^−1^) buffered with 0.1 M acetic acid buffer (pH 4.8) at 420 nm. One unit of enzyme activity (U) was defined as the amount of enzyme catalyzing the production of 1 μmol oxidized product per min. All measurements were performed in triplicate.

## RESULTS

### Laccase activity of *D. Squalens* in response to oxidative stress in mycelial interaction

When *D. squalens* (Ds) was co-cultured with *T. versicolor* (Tv) and *P.ostreatus* (Po) in Sc medium, the laccases activity was significantly increased in the interaction zones compared to single culture of *Ds* (Figure 1A), reaching 307.56 U/L in DsTv and 274.50U/L in DsPo, and enhanced changes in laccase activity were as high as 190.04-fold in DsTv and 212.93-fold in DsPo compared to the control. This strong laccase response matched previous findings for Ds during interactions with other species, but the extent of increase in laccase activity was more significant than that in other studies based on the different identity of the competitor and culture conditions (Dong et al., 2011; Kannaiyan et al., 2015). Besides, the result showed laccase activity in the interaction zones of DsTv and DsPo were all higher compared to other two single fungi Tv and Po (Figure 1A). We also identified laccase activity neighboring the interaction zone. Although laccase activity declined as distance increased from the interaction zone, it was still higher than that of the Ds single culture. The result showed that secondary metabolites produced by interspecific interaction could induce laccase activity. Since laccase production in the self-fungal interaction zone did not increase compared to the control (Figure S6), it is highly possible that interspecific antagonism contributed to the induction of laccase activity.

**Figure 1.**
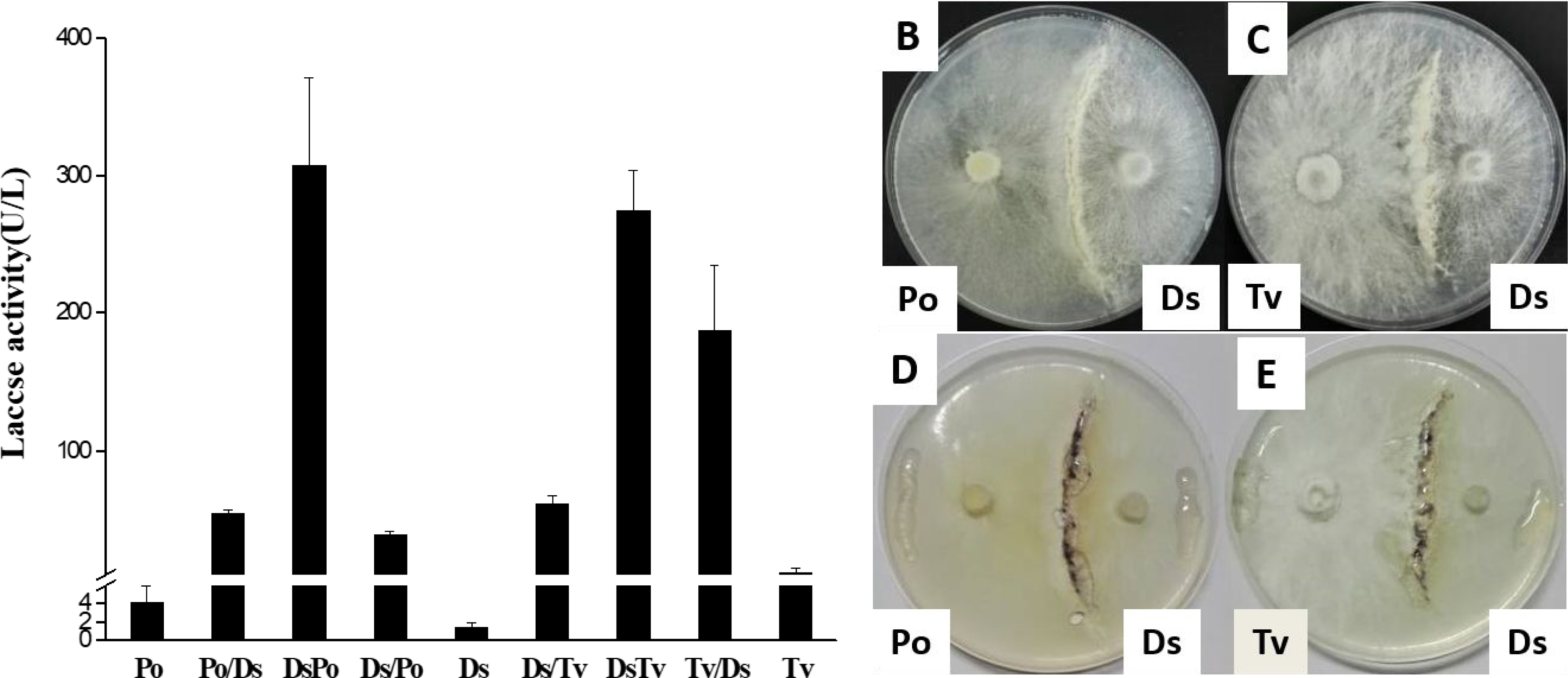
Interaction on Sc agar plate between *D. squalens* and other two competitors. (A) Laccase production of paired fungal interaction. Ds, Po, Tv: single culture of *D.squalens, P.ostreatus, T. versicolor.* DsTv: interaction zone of *D.squalens* and *T. versicolor*, DsPo: interaction zone of *D.squalens* and *P.ostreatus.* Ds/Tv: *D.squalens* region adjacent to the interaction zone of *D.squalens* and *T. versicolor.* Tv/Ds: *T. versicolor* region adjacent to the interaction zone of *D.squalens* and *T. versicolor.* Ds/Po: *D.squalens* region adjacent to the interaction zone of *D.squalens* and *P.ostreatus.* Po/Ds: *P.ostreatus* region adjacent to the interaction zone of *D.squalens* and *P.ostreatus.* Interaction zone of *D. squalens* and *P. ostreatus* (B) or *T. versicolor* (C) without nitroblue tetrazolium (NBT) staining. Purple coloration was observed in the interaction zone of *D. squalens* and *P. ostreatus* (D) or *T. versicolor* (E) with nitroblue tetrazolium (NBT) staining.

Research has suggested that overproduction of laccase during interspecific interaction is caused by oxidative stress, which involves a series of competitive and defensive mechanisms (Jaszek et al., 2006; Li et al., 2011; Yang et al., 2012). To determine oxidative stress during mycelial interaction, we used Nitroblue tetrazolium (NBT) to stain the mycelium in both interaction zones and single species (Figure 1D and E). Compared to the co-culture without NBT staining (Figure 1B and C), no color change was observed in the single species near the plate edge, but purple formazan precipitate formation was obviously observed in both interaction zones of DsTv and DsPo, which could indicate much higher accumulation of ROS during mycelial interactions.

### RNA-Seq analysis of gene expression

To gain a better understanding of the interaction mechanisms, RNA-Seq analysis was used to study the global gene expression of *Ds* when grown in pure culture and compared to the interaction zone of DsTv and DsPo. Illumina sequencing data was deposited in the NCBI SRA database with accession number SRR5328881. A minimum RPKM (Reads Per Kilo-base per Million) value threshold was set at two-fold to limit the false-positive detection of low-abundance genes. To validate the technical reproducibility of the RNA-seq experiment, we compared the expression values of three biological replicates. After sequencing, over 128.88 million raw reads and 6.57 billion raw bases were obtained in a single sequencing run (Table S5). The raw data was firstly filtered to obtain high-quality clean data to ensure the success of subsequent analysis. Finally, we obtained over 112.36 million clean reads and 5.48 clean bases (Table S5). According to the related data (Table S6), reads from Ds transcripts data sets mapped to 86.86% (average value) of its genes, while reads from DsTv and DsPo transcripts data sets mapped to 14.49% and 15.02% (average value) of Ds genes respectively. Although the mapping rate to Ds reads was low in two interactions, 98.23% of Ds genes could be detected in both interactions of DsTv and DsPo, and the total mapping rate was more than 60% when adding the mapping rate to Po and Tv reads. The raw data quality control (Figure S1-S3) indicated that the sequencing quality of three samples was high, and the sequencing data can meet the requirements of subsequent analysis.

### Analysis of differential expression genes (DEGs)

Given the results of the RNA-seq data, gene expression with a competitor-to-control ratio of RPKM higher than 2 or below 0.5 and FDR<0.05 were considered differentially expressed (Figure S4). The sequences of genes were searched against the NCBI non-redundant sequences (NR) database using BlastX and setting a cut-off E-value of 10^−5^. According to the result, a large number of Ds genes were found to be differentially expressed in the two interactions (DsPo and DsTv) compared to the control, and both two interactions comprised more up-regulated genes than down-regulated genes (Figure 2 and Table S1). Of these, 1085 genes were up-regulated and 873 were down-regulated in DsTv, while 1182 genes were up-regulated and 783 were down-regulated in DsPo. In addition, over half of genes were up- or down-regulated in both interactions, with 621 up-regulated genes and 441 down-regulated genes in both interactions.

**Figure 2.**
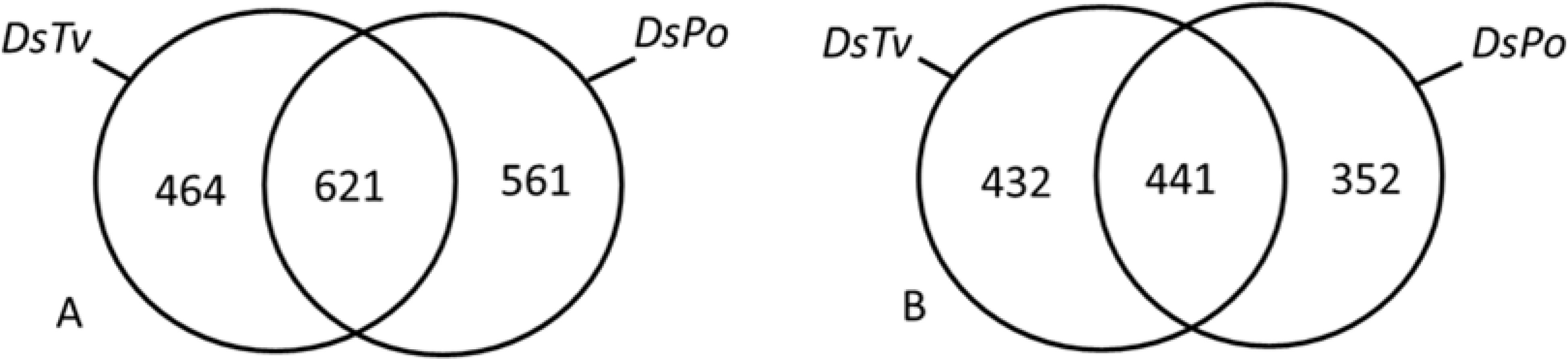
Venn diagrams of number of DEGs compared to control sample in the interaction with *T. versicolor* (DsTv) and *P. ostreatus* (DsPo). (A) Genes were up-regulated by at least 2-fold; (B) Genes were down-regulated by at least 2-fold.

Interestingly, 14 main oxidative stress-related and energy metabolism-related functional groups of DGEs annotated against NR database were found significantly up-regulated in both interactions DsTv and DsPo compared to the control (Figure 3). Of these functional groups, the largest one was cytochromes P450. This group composed of 52 DEGs in DsTv and 47 DEGs in DsPo, and most of which were up-regulated in two interactions. The function group of glycoside hydrolase also contained abundant DEGs, although the number of up-regulated DEGs in this group was less than down-regulated DEGs, the change folds of up-regulated DEGs was more significant in both interactions. For instance, one glycoside hydrolase-encoding gene (DICSQDRAFT_109102) could reach to 556.80-fold in DsTv and 961.61-fold in DsPo (Table S1). Moreover, other oxidative stress-related groups including alcohol oxidase, glutathione S-transferase, laccase, terpenoid synthase, manganese peroxidase, multidrug resistance-associated ABC transporter, alcohol dehydrogenase and FMN-linked oxidoreductase, and energy metabolism-related groups including 3-ketoacyl-CoA thiolase, S-adenosyl-L-methionine dependent methyltransferase, chitinase and ATPase were all composed of more highly up-regulated DEGs than down-regulated DEGs in DsTv and DsPo.

**Figure 3.**
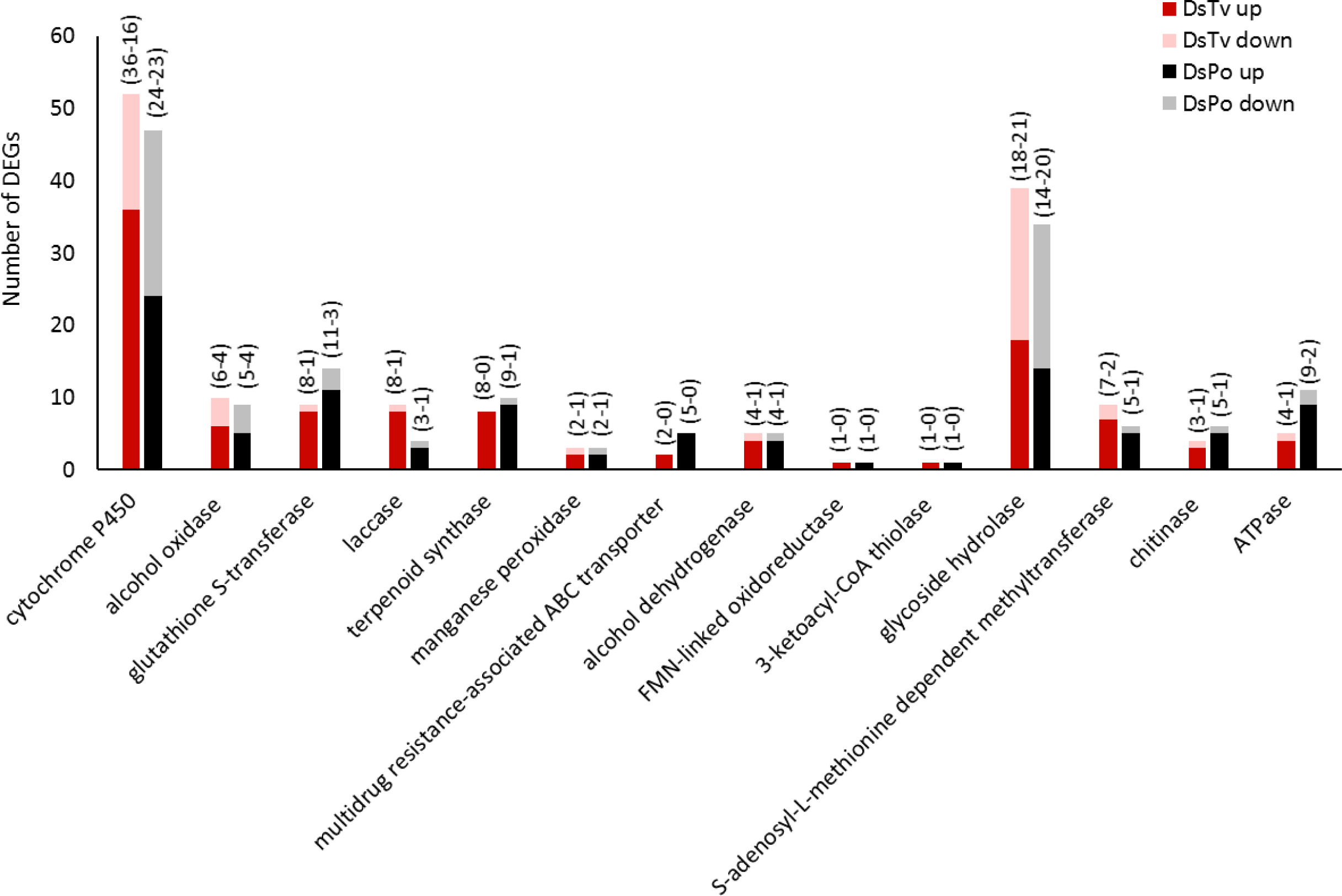
Expression patterns for 14 oxidative stress-related and energy metabolism-related functional groups of DEGs. For each functional group, the number of up- or down-regulated DEGs in the interaction of DsTv and DsPo compared to control are shown. Values between parentheses above each bar correspond to the number of up- and down-regulated DEGs in each functional group.

Importantly, glutathione S-transferase and laccase, which both function as detoxification and ROS scavengers (Yang et al., 2012), were remarkably induced in two interactions. For the glutathione S-transferase group, eight up-regulated and one down-regulated DEGs in DsTv, and 11 up-regulated and three down-regulated DEGs in DsPo, For the laccase group, eight and three up-regulated DEGs in DsTv and DsPo respectively, only one gene was down-regulated in two interactions. Among these, three laccase genes (Dslcc5, Dslcc10 and Dslcc11) were up-regulated in both interactions, and the up-regulated folds of Dslcc11 even could reach up to 115-fold in DsTv and 148-fold in DsPo (Table S7).

The 25 up-regulated genes with highest fold change under both conditions are presented in Table 2 (not including genes with unknown function). This analysis showed that most highly up-regulated genes encoded proteins which played a similar role in the interaction of DsTv or DsPo. Of these, many significantly up-regulated genes encoding proteins were involved in xenobiotic detoxification and generating or reducing ROS, including aldo/keto reductase, glutathione S-transferases, NADH:flavin oxidoreductase/NADH oxidase, short-chain dehydrogenases/ reductase, cytochrome P450 enzymes, the succinate dehydrogenase cytochrome b560 subunit, manganese peroxidase, aldehyde dehydrogenase and laccase. In addition, some gene encoding proteins were involved in an antagonistic mechanism of nutrient competition and antifungal properties, including glycoside hydrolase, glucanase, alpha/beta-hydrolase, GroEs-like protein and terpenoid synthases.

**Table 2.** The 25 up-regulated genes with highest fold change under both co-culture conditions by RNA-seq analysis

| Up-regulated genes in the interaction with *P. ostreatus* |  |  | Up-regulated genes in the interaction with *T. versicolor* |  |  |
| --- | --- | --- | --- | --- | --- |
| Gene_ID | Fold change | Putative function | Gene_ID | Fold change | Putative function |
| 18845230 | 2160.84 | oxidoreductase | 18845401 | 2057.23 | Non-Catalytic module family EXPN protein |
| 18845401 | 1525.30 | Non-Catalytic module family EXPN protein | 1884439 | 1157.90 | UbiA prenyltransferase |
| 18843716 | 976.27 | NAD(P)-binding protein | 18833825 | 556.79 | glycoside hydrolase |
| 18833825 | 961.61 | glycoside hydrolase | 18837948 | 328.53 | terpenoid synthase |
| 18837948 | 674.14 | terpenoid synthase | 18843669 | 316.56 | Nitroreductase |
| 18841174 | 622.22 | GroES-like protein | 18843554 | 288.29 | cytochrome P450 |
| 18842297 | 570.29 | Non-catalytic module family EXPN protein, partial | 18843877 | 203.59 | HAD-like protein, partial |
| 18844395 | 547.52 | UbiA prenyltransferase | 18842297 | 190.84 | Non-catalytic module family EXPN protein, partial |
| 18839563 | 343.20 | Aldo/keto reductase | 18844746 | 126.84 | amine oxidase |
| 18837949 | 316.56 | alpha/beta-hydrolase | 18833943 | 114.88 | laccase B |
| 18837763 | 255.86 | NADH:flavin oxidoreductase/NADH oxidase | 18839529 | 92.21 | succinate dehydrogenase cytochrome b560 subunit |
| 18842217 | 196.82 | Osmotin thaumatin-like protein | 18837704 | 87.02 | DUF323 domain-containing protein |
| 18843669 | 148.74 | Nitroreductase | 18842217 | 84.90 | Osmotin thaumatin-like protein |
| 18833943 | 148.38 | laccase B | 18837323 | 84.90 | arylalcohol dehydrogenase |
| 18834029 | 133.96 | carotenoid ester lipase | 18839046 | 84.55 | laccase |
| 18839183 | 127.52 | glutathione S-transferase C-terminal-like protein | 18845343 | 60.52 | secreted protein |
| 18840270 | 115.66 | Aldo/keto reductase | 18842241 | 58.78 | Manganese-dependent peroxidase |
| 18835851 | 112.08 | FAD-binding domain-containing protein | 18835851 | 52.59 | FAD-binding domain-containing protein |
| 18836474 | 104.35 | short-chain dehydrogenases/reductase | 18844326 | 52.57 | terpenoid synthase |
| 18836780 | 94.17 | barwin-like endoglucanase | 18844314 | 48.29 | GroES-like protein |
| 18834127 | 82.64 | NAD(P)-binding protein | 18833814 | 42.25 | cytochrome P450 |
| 18836407 | 78.55 | NAD(P)-binding protein | 18838354 | 42.20 | RTA1-domain-containing protein |
| 18842871 | 76.96 | NAD(P)-binding protein | 18842447 | 40.32 | GroES-like protein |
| 18844326 | 68.56 | terpenoid synthase | 18844012 | 40.10 | carbonic anhydrase, partial |
| 18837789 | 67.90 | Aldo/keto reductase | 18843657 | 39.62 | O-methyltransferase |

### GO and KEGG analysis

Gene Ontology (GO) terms enrichment analysis globally provides significantly enriched GO terms in DEGs between co-cultures and single culture. Using GO analysis, a total of 45 and 27 GO terms were enriched in DEGs with a p-value <0.05 in two interactions of DsTv and DsPo, respectively. Interestingly, we found the same number of GO terms (20 GO terms) with FDR <0.05 (FDR adjusted p-value<0.05) showed significant enrichment in DsTv and DsPo. Although few enriched GO terms were distinct in DsTv and DsPo, most were the same and had similar enrichment ratios (Figure 4). According to the result, only two main categories, Biological Process (BP) and Molecular Function (MF), contained significantly enriched GO terms in both interactions. Genes categorized into “Cellular Component” showed no significant enrichment (FDR>0.05, Table S3). Under each GO categories, a large number of DEGs were most significantly (FDR<0.001) enriched in categories of “oxidation-reduction process,” “oxidoreductase activity,” “heme binding,” “oxidoreductase activity with incorporation or reduction of molecular oxygen,” “coenzyme binding,” “cofactor binding,” “iron ion binding” and “monooxygenase activity” in both interactions, which suggested that the response to stressful conditions was induced during interspecific interaction. We noticed that the highest percentage of DEGs were exclusively enriched in DsPo categories of “glutathione conjugation reaction” and “glutathione transferase activity,”, while several DEGs were exclusively enriched in DsTv categories of “carbohydrate metabolic process,” “ion binding” and “flavin adenine dinucleotide binding.” As more DEGs of significantly enriched GO terms were up-regulated, these GO annotations also indicated that DEGs encoding proteins with antioxidant activity and detoxification properties were more active during the mycelial interaction.

**Figure 4.**
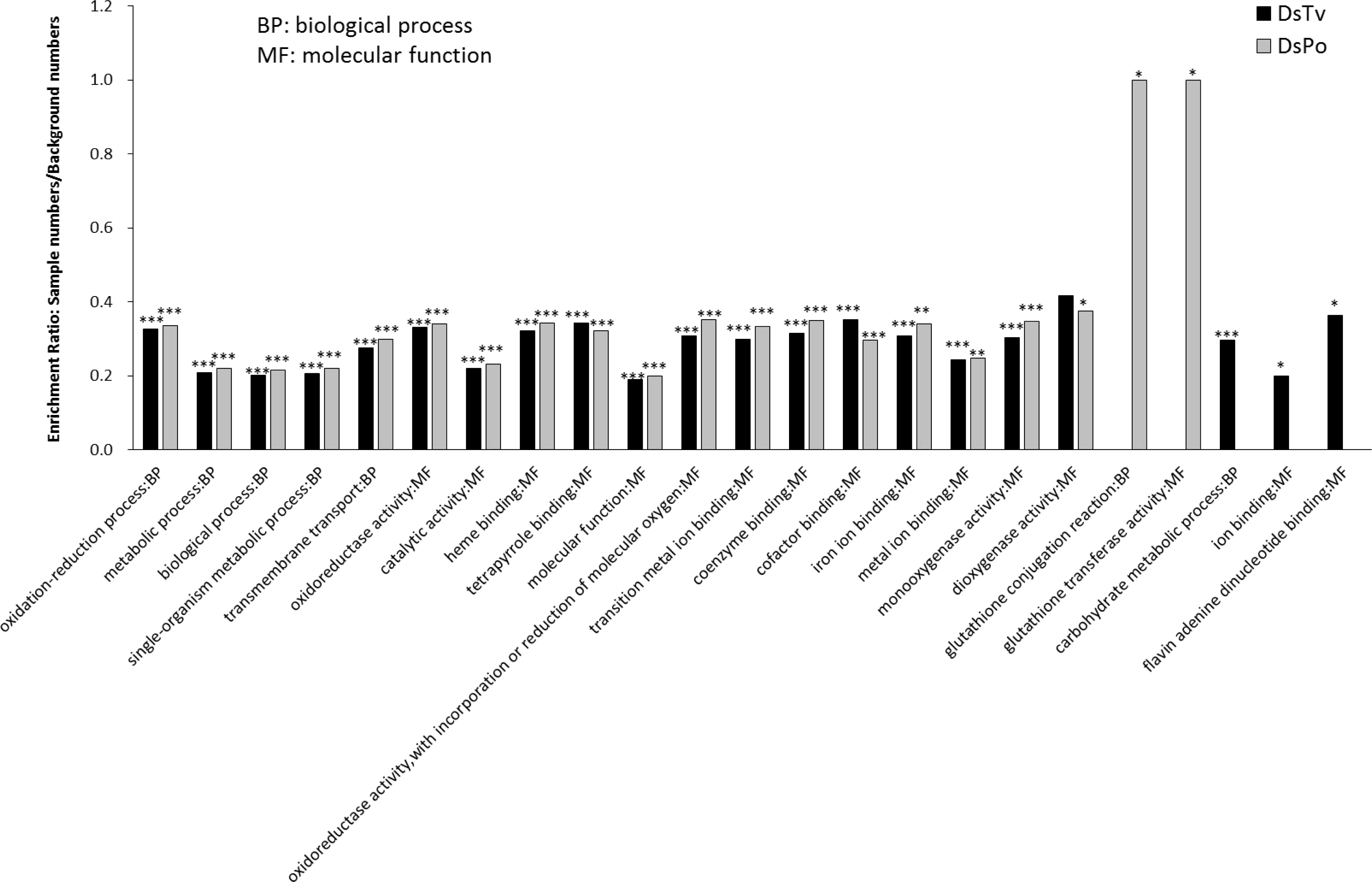
GO terms enrichment analysis on both interactions compared to control. (*** FDR<0.001, **FDR<0.01, * FDR<0.05)

DEGs usually cooperate to execute their biological functions, and KEGG pathway analysis is widely used to further understand the biological functions of genes. KEGG pathway enrichment analysis was used to identify significantly enriched metabolic pathways or signal transduction pathways in DEGs. Contrary to GO analysis, the DEGs of two interactions, DsTv and DsPo, showed distinct enrichment in KEGG pathways (Table S4). A total of 25 and 22 pathways enriched in DEGs with a p-value <0.05 were selected from the interaction of DsTv and DsPo, respectively (Figure 5). We noticed that the pathways of “Drug metabolism-cytochrome P450”, “Glutathione metabolism,” “Metabolism of xenobiotics by cytochrome P450,” “Phenylpropanoid biosynthesis,” “chloroalkane and chloroalkene degradation,” “terpenoid backbone biosynthesis” and “Synthesis and degradation of ketone bodies” were significantly enriched in DEGs of both interactions. Moreover, pathways of “Aminobenzoate degradation,” “Dioxin degradation,” “Polycyclic aromatic hydrocarbon degradation” and “Riboflavin metabolism” were only significantly enriched in DEGs of DsPo, while “Bisphenol degradation,” and “Chloroalkane and chloroalkene degradation” were only significantly enriched in DEGs of DsTv. The KEGG analysis also showed that most KEGG pathways are mainly involved in xenobiotics metabolism and biodegradation, which may also function as defense responses to toxic substances and biological attacks.

**Figure 5.**
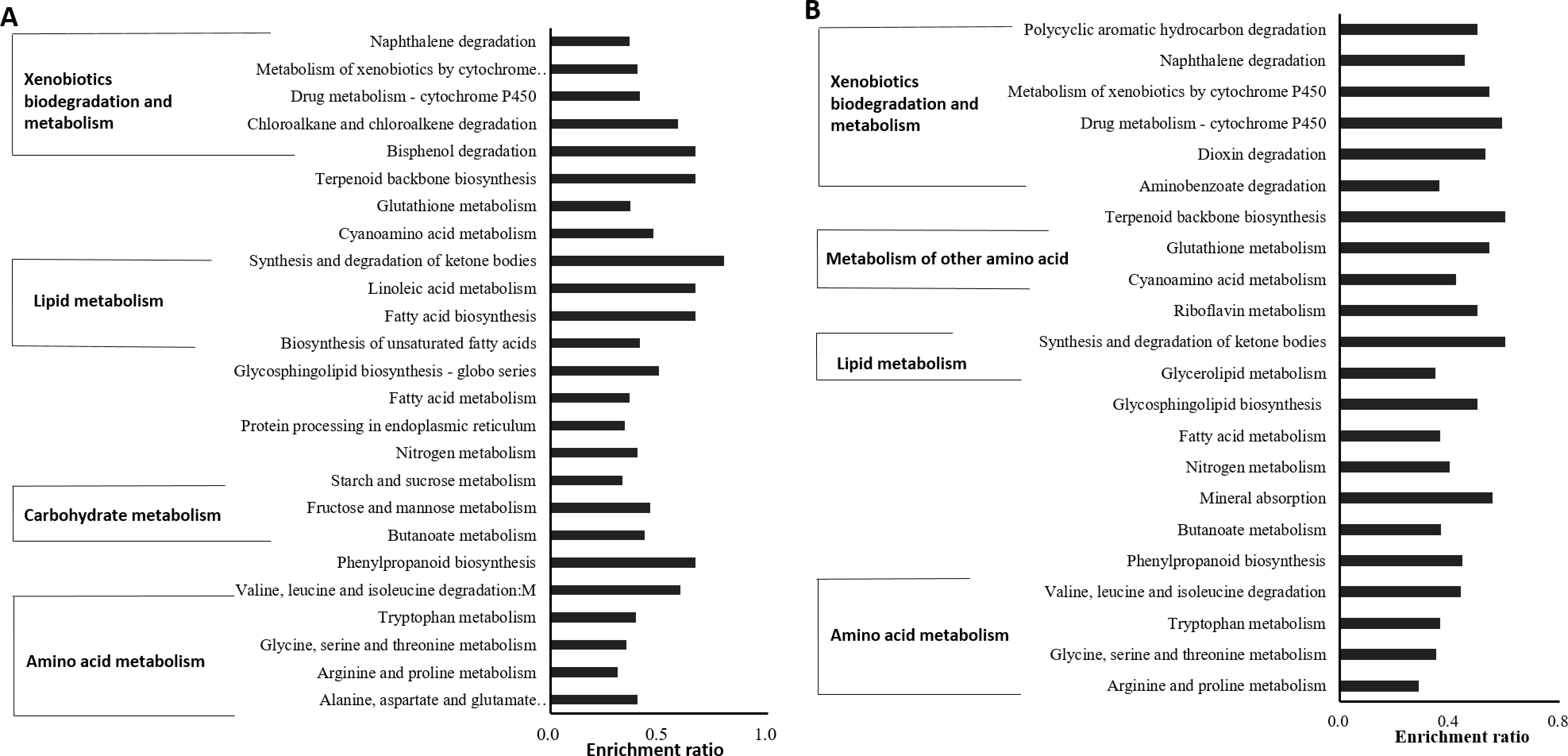
KEGG pathway enrichment analysis in *D. Squalens* under the interaction with (A) *T. versicolor* and (B) *P ostreatus* compared to the control. (p-value<0.05)

Apart from a defensive function, nutrient competition also existed for mycelium growth and development during this interspecific interaction (Boddy, 2000; Heilmann-Clausen and Boddy, 2005; Wells and Boddy, 2002). KEGG analysis indicated that some significantly enriched pathways were involved in energy production and metabolism of both interactions. The up-regulated “Nitrogen metabolism” and amino acid metabolism such as “Arginine and proline metabolism” and “Glycine, serine and threonine metabolism” were indispensable for fungal growth when interacting with its competitors. Carbohydrate metabolic pathways such as “Fructose and mannose metabolism” and “Starch and sucrose metabolism” and amino acid metabolic pathways such as “Alanine, aspartate and glutamate metabolism” were only significantly enriched in the DEGs of DsTv, suggesting that the antagonism mechanism of Ds in nutrition competition depended on the type of interaction partner. As oxidative phosphorylation pathway was intensified in both PoDs and PoTv (Figure S5), which also implied the ATP production and metabolism was enhanced for intensive nutrient competition.

### Validation of the DEGs results by qRT-PCR analysis

Based on the RNA-Seq analysis, target genes were selected for quantitative real-time PCR to validate the gene expression profile. According to the RNA-seq result, we selected 10 genes that were all up-regulated in both interactions and had functional annotation involved in ROS production and regulation, including two laccases (Dslccll and Dslcc5), NAD(P)-binding protein (DICSQDRAFT_70438), aldo/keto reductase (DICSQDRAFT_172438), glutathione S-transferase (DICSQDRAFT_161976), GroES-like protein (DICSQDRAFT_182440), NADH:flavin oxidoreductase (DICSQDRAFT_157369), cytochrome P450 monooxygenase (DICSQDRAFT_55106) and acetyl-CoA synthetase-like protein (DICSQDRAFT_89216) and Dyp-type peroxidase (DyP1). We also selected six down-regulated genes that all represented large functional groups in both interactions, including cytochrome P450 (DICSQDRAFT_130358), FAD/NAD(P)-binding domain-containing protein (DICSQDRAFT_172084), NAD(P)-binding protein (DICSQDRAFT_58923), Aldo/keto reductase (DICSQDRAFT_167996), Clavaminate synthase-like protein (DICSQDRAFT_108315) and glycoside hydrolase (DICSQDRAFT_149582).

The relative expression values of a total of 16 genes in two interactions of DsTv and DsPo provided by qRT-PCR were set compared to the control sample (Figure 6). The qRT-PCR results of 10 up-regulated and 6 down-regulated genes were all in accordance with the RNA-seq data. Of these five genes, DICSQDRAFT_70438, DICSQDRAFT_172438, DICSQDRAFT_157369, DyP1 and Dslcc11 were more highly up-regulated and one gene DICSQDRAFT_130358 was more down-regulated than other genes. We also compared the relative expression of each gene determined by qRT-PCR to values from the RNA-Seq analysis. The up- or down-regulated folds of most genes in qRT-PCR and RNA-Seq were of the similar orders of magnitude, but there were distinct differences between higher-value RNA-Seq and qRT-PCR data (Figure S7). The results suggested that the RNA-Seq data and DEGs analysis during the interaction of DsTv and DsPo were reliable.

**Figure 6.**
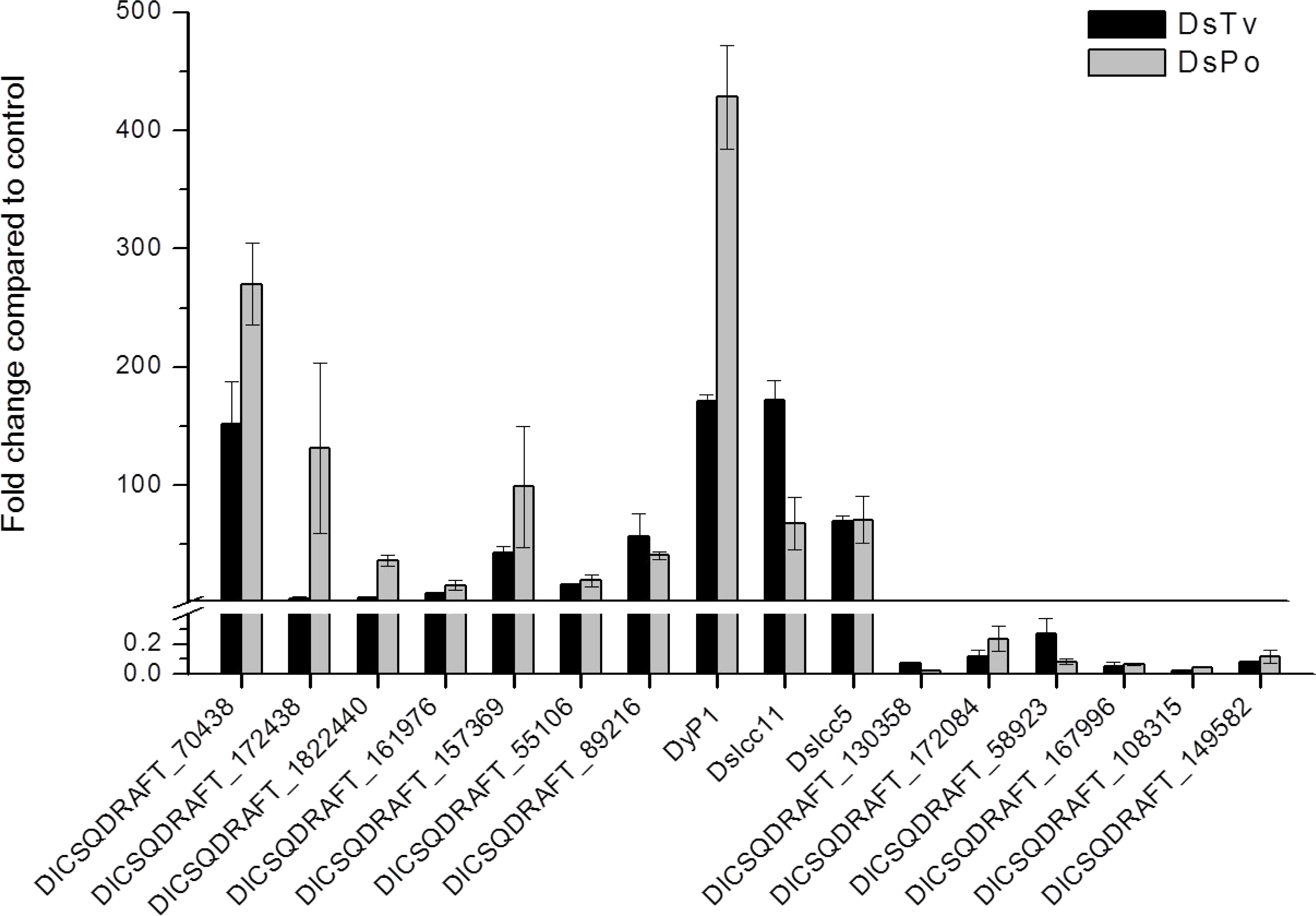
Validation of up-and down-regulated genes expression during both interactions compared to control by quantitative RT-PCR. Expression levels of 16 genes in *D. squalens* were set as control and normalized to 1. Black horizontal line corresponds to level equal to control. Values are averaged from three biological replicates.

## DISCUSSION

In this study, the potential of Ds to enhance laccase production by co-culturing with Tv or Po was identified. Notably, laccase activity were remarkably increased in DsTv and DsPo compared to the control. As the expression of two major laccase genes Dslcc11 and Dslcc5 were also significantly up-regulated based on RNA-seq analysis (Table S7), suggesting that Dslcc11 and Dslcc5 probably play key roles in laccase activity induction. Although many studies have revealed that interspecific fungal interactions contribute to the increase in laccase activity (Baldrian, 2004; Chi et al., 2007; Ferreira Gregorio et al., 2006; Flores et al., 2009; Hiscox et al., 2010; Kuhar et al., 2015; Wei et al., 2010), the mechanism of laccase production caused by mycelial interactions remains elusive. As no research has included analysis of the differential expression of genes of Ds when Ds interacted with other fungi, these two interactions of DsTv and DsPo were chosen for further study of laccase in response to interspecific antagonism.

The RNA-seq data showed remarkable changes in Ds genetic expression, with 13.64% of the genes in DsTv and 15.90% of the genes in DsPo differentially expressed, similar to the gene expression change of other fungal interactions (Arfi et al., 2013). Moreover, the number of up-regulated DEGs during the interactions was also similar to the number of down-regulated DEGs, but the up-regulated folds of most DEGs were generally higher than the down-regulated folds compared to the control (Table S1). The DEGs analysis by RNA-Seq was confirmed to be reliable by performing qRT-PCR analysis of the expression of a 16-gene set.

It has been reported that interspecific interaction could induce an oxidative stress response by accumulation of ROS (Iakovlev et al., 2004; Silar, 2005). Toxic ROS can lead to cellular death by damaging DNA and proteins and triggering lipid peroxidation (Garcia-Sanchez et al., 2012; Ujor et al., 2012), likely inducing the cellular functions of detoxification, defense and repair (Ujor et al., 2012). In our study, oxidative stress/protective response may be active in the interaction zone owing to the up-regulation of many DEGs encoding functional groups related to ROS generation, including alcohol oxidases, which can produce H_2_O_2_ to support Fenton reactions (Daniel et al., 2007), and Cytochrome P450 enzymes, which are involved in xenobiotic metabolism at the expense of increasing ROS during the catalytic cycle process (Gonzalez, 2005).

To resist damage due to ROS, detoxification and defense response could be stimulated to mediate oxidative stress, which is associated with a series of second metabolites secreted from fungi (Evans et al., 2008; Jonkers et al., 2012). Interestingly, in our data, the putative functional groups encoded by a large number of DEGs appear to play a similar role in removing ROS and detoxifying xenobiotics (Table S2). For instance, aldo/keto reductase are responsible for the reductive metabolism of biogenic and xenobiotic materials to primary and secondary alcohols, leading to their detoxification (Mazhawidza et al., 2014). Aldehyde dehydrogenase belongs to detoxification enzymes due to its role in the metabolism of intermediates and exogenous aldehydes and consequently protect cell against oxidative stress (Chen et al., 2014). Glutathione S-transferases are involved in xenobiotic metabolism and cellular detoxification during oxidative stress response (Mariani et al., 2008). Lignin-modifying enzymes including laccase and manganese peroxidase are also important for detoxification and resistance against oxidative stress (Aust, 2004; Jaszek et al., 2006). Based on our RNA-seq analysis, the laccase group was highly up-regulated in both interactions, which is consistent with strongly increased activity of laccase during interactions. Although the previous study of the *Pycnoporus coccineus* also showed that most highly up-regulated genes in *P. coccineus* under interaction with *Botrytis cinerea* and *Coniophora puteana* were annotated as aldo/keto reductases, glutathione S-transferases and cytochrome P450 enzymes (Arfi et al., 2013), there are still some differences compared with the result in our study. The most important one is that no differential expressed laccase gene was identified in this paper, which cannot be related to the overproduction of laccase in white-rot fungi under interaction.

As laccases are considered to play a key role in the antagonistic mechanisms, their overexpression also implies a strong competition of nutrients in the mycelial interaction zone. When two different fungi competed for resource and territory, a series of antifungal metabolites were released to protect against competitors and assist in the acquisition of nutrients (Wang M, 2013), such as toxic phenolic compounds and terpenoids, both of which can induce laccase production (Ferreira Gregorio et al., 2006). The up-regulation of DEGs encoding terpenoid synthases in RNA-seq data indicated the secretion of terpenoids, organic metabolites with antifungal properties that are involved in the defense responses of fungi (Evans et al., 2008; Yap et al., 2015). Phenolic compounds can directly generate the accumulation of ROS and consequently produce oxidative stress (Cruz-Ortega R, 2007; Garcia-Sanchez et al., 2012). The role of laccase may be to catalyze the monoelectronic oxidation of phenols released by its competitors to the corresponding radical to detoxify or repress ROS (Ferreira Gregorio et al., 2006). The phenolic substrates are also used as laccase mediators, whose oxidized radicals lead to the formation of new dimers, oligomers and polymers via binding to non-laccase substrates (Jeon et al., 2012; Riva, 2006). Moreover, we found abundant genes encoding carbohydrate-active enzymes (CAZy) were upregulated in both interaction zones, including glycoside hydrolase and glucanase, which also reflected intensive competition in territory and nutrient uptake when different fungi species confronted with each other.

The GO and KEGG analysis of the DEGs also suggested that oxidative stress exists in the interaction zone. According to the significant enriched GO categories of DEGs, most of DEGs with a role in the response to stressful conditions were up-regulated during the interaction of DsTv or DsPo. The remarkably up-regulated genes mainly encode NAD(P)-binding protein (DICSQDRAFT_70438), terpenoid synthase (DICSQDRAFT_159719), GroES-like protein (DICSQDRAFT_182440), aldo/keto reductase (DICSQDRAFT_17243 8), NADH:flavin oxidoreductase (DICSQDRAFT_ 157369), laccase B (Dslcc11), glutathione S-transferase C-terminal-like protein (DICSQDRAFT_170658), and FAD-binding domain-containing protein (DICSQDRAFT_139420), these proteins exactly belong to the significantly enriched GO categories of “oxidation-reduction process,” “metabolic process,” “single-organism metabolic process,” “oxidoreductase activity,” “catalytic activity,” “molecular function,” “transition metal ion binding,” “coenzyme binding,” “cofactor binding,” “metal ion binding,” and “monooxygenase activity,” which likely indicates that intracellular molecule binding, xenobiotics metabolism and oxidative stress resistance were more active in Ds when it interacted with other fungi. This was in agreement with the previous report that laccase was involved in the response to oxidative stress (Jaszek et al., 2006; Piscitelli et al., 2011). Based on the analysis of KEGG pathways, enhanced carbohydrate metabolism in cells was associated with the activation of defense and antioxidative stress responses during the interspecific interaction (Shetty et al., 2007), and DEGs involved in carbohydrate and nitrogen metabolism could be related to nutrition competition, in accordance with the fungal ability to take up carbon from the competitor’s mycelium in the interaction zone (Eyre et al., 2010; Wells and Boddy, 2002). In addition, the up-regulation of fatty acid metabolism and amino acid metabolism with energy production was also correlated with the combative mechanism and laccase induction.

In conclusion, our results provide insights into antagonistic mechanisms of interspecific interaction between white rot fungi by analyzing differential gene expression and revealed that laccase induction was probably the result of oxidative stress/protective responses and nutrient competition with opposing competitors since many oxidative stress-resistant genes, antifungal genes and carbohydrate-active enzymes-encoding genes were significantly up-regulated. Importantly, we presented three highly up-regulated laccase genes (Dslcc5, Dslcc10 and Dslcc11) that can be examined for their transcription factor in future experiments.

## Acknowledgements

This study was funded by the National Natural Foundation of China (31400063), Fundamental Research Funds for the Central Universities (No. XDJK2011B009; XDJK2017B030), Research Funds of Scientific Platform and Base Construction (No. cstc2014pt-sy0017), Chongqing Research Program of Social undertaking and livelihood security (No. cstc2016shmszx1176) and The Recruitment Program for Foreign Experts (No. WQ20125500073), and The State Bureau of Forestry 948 project (No. 2015-4-42).

## Author contributions

Z.Z. and N.L. performed the experiments and wrote the manuscript, and Y.I., B.H. and F.L. designed the experiments and revised the manuscript. All authors discussed and commented on the manuscript.

## Additional Informantion

Competing financial interests: The authors declare no competing financial interests.

## References

Arfi Y, Levasseur A, Record E, 2013. Differential Gene Expression in Pycnoporus coccineus during Interspecific Mycelial Interactions with Different Competitors. Applied and environmental microbiology. 79, p. 6626–36.

Aust SD, Detoxification and Metabolism of Chemicals by White-Rot Fungi. Book Detoxification and Metabolism of Chemicals by White-Rot Fungi, 2004.

Baldrian P, 2004. Increase of laccase activity during interspecific interactions of white-rot fungi. FEMS microbiology ecology. 50, 245–53.

Boddy L, 2000. Interspecific combative interactions between wood-decaying basidiomycetes. FEMS Microbiol Ecol. 31, 185–94.

Chen CH, Ferreira JCB, Gross ER, Mochly-Rosen D, 2014. Targeting aldehyde dehydrogenase 2:new therapeutic opportunities. Physiol Rev. 94, 1–34.

Chi Y, Hatakka A, Maijala P, 2007. Can co-culturing of two white-rot fungi increase lignin degradation and the production of lignin-degrading enzymes? International Biodeterioration & Biodegradation. 59, 32–9.

Cho NS, Wilkolazka AJ, Staszczak M, Cho HY, Ohga S, 2009. The Role of Laccase from White Rot Fungi to Stress Conditions. Journal-faculty of agriculture Kyushu University. 54, 81–3.

Cruz-Ortega R L-NA, Anaya AL, 2007. Allelochemical Stress Can Trigger Oxidative Damage in Receptor Plants. Plant Signal Behav. 2 269–70.

Daniel G, Volc J, Filonova L, Plihal O, Kubatova E, Halada P, 2007. Characteristics of Gloeophyllum trabeum alcohol oxidase, an extracellular source of H2O2 in brown rot decay of wood. Applied and environmental microbiology. 73, 6241–53.

Dong Y-C, Wang W, Hu Z-C, Fu M-L, Chen Q-H, 2011. The synergistic effect on production of lignin-modifying enzymes through submerged co-cultivation of Phlebia radiata, Dichomitus squalens and Ceriporiopsis subvermispora using agricultural residues. Bioprocess and Biosystems Engineering. 35, 751–60.

Eggert C, Temp U, Eriksson KE, 1996. The ligninolytic system of the white rot fungus Pycnoporus cinnabarinus: purification and characterization of the laccase. Applied & Environmental Microbiology. 62, 1151.

Eisenman HC, Casadevall A, 2012. Synthesis and assembly of fungal melanin. Applied microbiology and biotechnology. 93, 931–40.

Eisenman HC, Mues M, Weber SE, Frases S, Chaskes S, Gerfen G, et al., 2007. Cryptococcus neoformans laccase catalyses melanin synthesis from both D- and L-DOPA. Microbiology. 153, 3954–62.

El Ariebi N, Hiscox J, Scriven SA, Müller CT, Boddy L, 2016. Production and effects of volatile organic compounds during interspecific interactions. Fungal Ecology. 20, 144–54.

Evans JA, Eyre CA, Rogers HJ, Boddy L, Müller CT, 2008. Changes in volatile production during interspecific interactions between four wood rotting fungi growing in artificial media. Fungal Ecology. 57–68.

Eyre C, Muftah W, Hiscox J, Hunt J, Kille P, Boddy L, et al., 2010. Microarray analysis of differential gene expression elicited in Trametes versicolor during interspecific mycelial interactions. Fungal biology. 114, 646–60.

Ferreira Gregorio AP, Da Silva IR, Sedarati MR, Hedger JN, 2006. Changes in production of lignin degrading enzymes during interactions between mycelia of the tropical decomposer basidiomycetes Marasmiellus troyanus and Marasmius pallescens. Mycological research. 110, 161–8.

Flores C, Vidal C, Trejo-Hernandez MR, Galindo E, Serrano-Carreon L, 2009. Selection of Trichoderma strains capable of increasing laccase production by Pleurotus ostreatus and Agaricus bisporus in dual cultures. Journal of applied microbiology. 106, 249–57.

Floudas D, Binder M, Riley R, Barry K, Blanchette RA, Henrissat B, et al., 2012. The Paleozoic origin of enzymatic lignin decomposition reconstructed from 31 fungal genomes. Science. 336, 1715.

Flundas D, Hibbett DS, 2012. The Paleozoic Origin of Enzymatic Lignin Decomposition Reconstructed from 31 Fungal Genomes. Science. 336, 1715.

Garcia-Sanchez M, Garrido I, Casimiro Ide J, Casero PJ, Espinosa F, Garcia-Romera I, et al., 2012. Defence response of tomato seedlings to oxidative stress induced by phenolic compounds from dry olive mill residue. Chemosphere. 89, 708–16.

Gonzalez FJ, 2005. Role of cytochromes P450 in chemical toxicity and oxidative stress: studies with CYP2E1. Mutation research. 569, 101–10.

Heilmann-Clausen J, Boddy L, 2005. Inhibition and stimulation effects in communities of wood decay fungi: exudates from colonized wood influence growth by other species. Microbial ecology. 49, 399–406.

Hiscox J, Baldrian P, Rogers HJ, Boddy L, 2010. Changes in oxidative enzyme activity during interspecific mycelial interactions involving the white-rot fungus Trametes versicolor. Fungal genetics and biology: FG & B. 47, 562–71.

Iakovlev A, Olson A, Elfstrand M, Stenlid J, 2004. Differential gene expression during interactions between Heterobasidion annosum and Physisporinus sanguinolentus. FEMS Microbiol Lett. 241, 79–85.

Jaszek M, Grzywnowicz K, Malarczyk E, Leonowicz A, 2006. Enhanced extracellular laccase activity as a part of the response system of white rot fungi: Trametes versicolor and Abortiporus biennis to paraquat-caused oxidative stress conditions. Pesticide Biochemistry and Physiology. 85, 147–54.

Jeon JR, Baldrian P, Murugesan K, Chang YS, 2012. Laccase-catalysed oxidations of naturally occurring phenols: from in vivo biosynthetic pathways to green synthetic applications. Microbial biotechnology. 5, 318–32.

Jonkers W, Rodriguez Estrada AE, Lee K, Breakspear A, May G, Kistler HC, 2012. Metabolome and transcriptome of the interaction between Ustilago maydis and Fusarium verticillioides in vitro. Applied and environmental microbiology. 78, 3656–67.

Kannaiyan R, Mahinpey N, Kostenko V, Martinuzzi RJ, 2015. Nutrient media optimization for simultaneous enhancement of the laccase and peroxidases production by coculture of Dichomitus squalens and Ceriporiopsis subvermispora. Biotechnology and applied biochemistry. 62, 173–85.

Kuhar F, Castiglia V, Levin L, 2015. Enhancement of laccase production and malachite green decolorization by co-culturing Ganoderma lucidum and Trametes versicolor in solid-state fermentation. International Biodeterioration & Biodegradation. 104, 238–43.

Lara OT, Riveros RH, Aguirre J, 2003. Reactive oxygen species generated by microbial NADPH oxidase NoxA regulate sexual development in Aspergillus nidulans. Molecular microbiology. 50, 1241–55.

Li B, Dewey CN, 2011. RSEM: accurate transcript quantification from RNA-Seq data with or without a reference genome. BMC Bioinformatics. 12, 323.

Li Q, Bai Z, O’Donnell A, Harvey LM, Hoskisson PA, McNeil B, 2011. Oxidative stress in fungal fermentation processes: the roles of alternative respiration. Biotechnology letters. 33, 457–67.

Livak KJ, Schmittgen TD, 2001. Analysis of relative gene expression data using real-time quantitative PCR and the 2(-Delta Delta C(T)) Method. Methods. 25, 402–8.

Mariani D, Mathias CJ, da Silva CG, Herdeiro Rda S, Pereira R, Panek AD, et al., 2008. Involvement of glutathione transferases, Gtt1and Gtt2, with oxidative stress response generated by H2O2 during growth of Saccharomyces cerevisiae. Redox report: communications in free radical research. 13, 246–54.

Mayer AM, Staples RC, 2002. Laccase: new functions for an old enzyme. Phytochemistry. 601, 551–65.

Mazhawidza W, Banda Y, Rajendran N, 2014. Bioinformatic Identification of Aldo-Keto Reductase from Newly Isolated Arthrobacter nicotianae strain PR and Its Phylogenetic Analysis Among Soil Bacteria. The Open Access Journal of Science and Technology. 2, 1–7.

Nosanchuk JD CA, 2003. The contribution of melanin to microbial pathogenesis. Cellular Microbiology. 5, 203–23.

Peiris D, Dunn W, Brown M, Kell D, Roy I, Hedger J, 2007. Metabolite profiles of interacting mycelial fronts differ for pairings of the wood decay basidiomycete fungus, Stereum hirsutum with its competitors Coprinus micaceus and Coprinus disseminatus. Metabolomics. 4, 52–62.

Piscitelli A, Giardina P, Lettera V, Pezzella, C., Sannia G, Faraco V, 2011. Induction and Transcriptional Regulation of Laccases in Fungi. Current Genomics. 12, 104–12.

Riva S, 2006. Laccases: blue enzymes for green chemistry. Trends in biotechnology. 24, 219–26.

Robinson MD, McCarthy DJ, Smyth GK, 2010. edgeR: a Bioconductor package for differential expression analysis of digital gene expression data. BIOINFORMATICS. 26, 139–40.

Shetty NP, Mehrabi R, Lutken H, Haldrup A, Kema GH, Collinge DB, et al., 2007. Role of hydrogen peroxide during the interaction between the hemibiotrophic fungal pathogen Septoria tritici and wheat. The New phytologist. 174, 637–47.

Silar P, 2005. Peroxide accumulation and cell death in filamentous fungi induced by contact with a contestant. Mycological research. 109, 137–49.

Ujor VC, Monti M, Peiris DG, Clements MO, Hedger JN, 2012. The mycelial response of the white-rot fungus, Schizophyllum commune to the biocontrol agent, Trichoderma viride. Fungal biology. 116, 332–41.

Wang M ea, 2013. Transcriptome and proteome exploration to provide a resource for the study of Agrocybe aegerita. PloS one. 8, e56686.

Wei F, Hong Y, Liu J, Yuan J, Fang W, Peng H, et al., 2010. Gongronella sp induces overproduction of laccase in Panus rudis. Journal of basic microbiology. 50, 98–103.

Wells JM, Boddy L, 2002. Interspecific carbon exchange and cost of interactions between basidiomycete mycelia in soil and wood. Functional ecology. 16, 153–61.

Xie C, Mao X, Huang J, Ding Y, Wu J, Dong S, et al., 2011. KOBAS 2.0: a web server for annotation and identification of enriched pathways and diseases. Nucleic acids research. 39, W316–22.

Yang Y, Fan F, Zhuo R, Ma F, Gong Y, Wan X, et al., 2012. Expression of the laccase gene from a white rot fungus in Pichia pastoris can enhance the resistance of this yeast to H2O2-mediated oxidative stress by stimulating the glutathione-based antioxidative system. Applied and environmental microbiology. 78, 5845–54.

Yap HY, Chooi YH, Fung SY, Ng ST, Tan CS, Tan NH, 2015. Transcriptome Analysis Revealed Highly Expressed Genes Encoding Secondary Metabolite Pathways and Small Cysteine-Rich Proteins in the Sclerotium of Lignosus rhinocerotis. PloS one. 10, e0143549.

Zhu X, Williamson PR, 2004. Role of laccase in the biology and virulence of Cryptococcus neoformans. FEMS yeast research. 5, 1–10.

